# On the robustness of graph-based clustering to random network alterations

**DOI:** 10.1101/2020.04.24.059758

**Authors:** R. Greg Stacey, Michael A. Skinnider, Leonard J. Foster

## Abstract

Biological functions emerge from complex and dynamic networks of protein-protein interactions. Because these protein-protein interaction networks, or interactomes, represent pairwise connections within a hierarchically organized system, it is often useful to identify higher-order associations embedded within them, such as multi-member protein-complexes. Graph-based clustering techniques are widely used to accomplish this goal, and dozens of field-specific and general clustering algorithms exist. However, interactomes can be prone to errors, especially interactomes that infer interactions using high-throughput biochemical assays. Therefore, robustness to network-level variability is an important criterion for any clustering algorithm that aims to generate robust, reproducible clusters. Here, we tested the robustness of a range of graph-based clustering algorithms in the presence of network-level noise, including algorithms common across domains and those specific to protein networks. We found that the results of all clustering algorithms measured were profoundly sensitive to injected network noise.

Randomly rewiring 1% of network edges yielded up to a 57% change in clustering results, indicating that clustering markedly amplified network-level noise. However, the impact of network noise on individual clusters was not uniform. We found that some clusters were consistently robust to injected network noise while others were not. Therefore, we developed the *clust.perturb* R package and Shiny web application, which measures the reproducibility of clusters by randomly perturbing the network. We show that *clust.perturb* results are predictive of real-world cluster stability: poorly reproducible clusters as identified by *clust.perturb* are significantly less likely to be reclustered across experiments. We conclude that quantifying the robustness of a cluster to network noise, as implemented in *clust.perturb*, provides a powerful tool for ranking the reproducibility of clusters, and separating stable protein complexes from spurious associations.

## INTRODUCTION

Networks are an important tool for representing the connections within a system, such as the agglomeration of proteins into complexes. Because these networks are composed of a list of connections (edges) between members (nodes), and do not explicitly detail higher-order associations, it can be useful to infer higher-order arrangements from the network. This task, called community detection or graph-based clustering, is ubiquitous across fields, and is especially important in biology, where the function of a biological macromolecule such as a protein is often mediated by its interacting partners within the network.

However, noise in networks can complicate clustering. This is especially true in biological networks constructed from high-throughput experiments, such as protein-protein interaction networks (“interactomes”) where more than half of the expected network edges may vary from experiment-to-experiment, either because of errors in network reconstruction or changes in experimental conditions (Stacey et al. 2018). Complicating this issue is the fact that it can be surprisingly ambiguous to measure differences between sets of clusters, likely because metrics for this purpose make choices about how penalize false positives (incorrectly merging clusters) versus false negatives (incorrectly separating clusters). This choice of weighting can mean popular metrics display biases and other non-intuitive behaviour (Gates et al. 2017), and papers using these metrics can include in-depth discussions of their behaviour (Nepusz, Yu, and Paccanaro 2012). This may explain why there is some discrepancy in the literature regarding the degree of noise-sensitivity when clustering biological networks, with some papers finding noise-sensitivity (Sloutsky et al. 2013) and others not (Brohée and van Helden 2006; Vlasblom and Wodak 2009; Freytag et al. 2018).

The aim of the present study study was to quantify the relationship between the level of network noise and cluster reproducibility. Our analysis focuses primarily on protein-protein interaction networks, but we reproduce our central findings in other types of networks. We first identified an unbiased cluster set similarity metric that behaved intuitively. With this in hand, we then randomly altered unweighted networks to varying degrees and measured the effects on the derived clusters. In order to arrive at general findings, we analyzed a literature-curated protein-protein interaction network (Giurgiu et al. 2019; Wishart et al. 2018), a second biological network (Wishart et al., 2018), a representative social network (Leskovec and Mcauley 2012; Yin et al. 2017), and 28 protein-protein interaction networks derived from four datasets generated by our group (Scott et al. 2017, 2015; Kristensen, Gsponer, and Foster 2012; Kerr et al., n.d.). We quantified the robustness of clusters obtained from five clustering algorithms that are widely used in a number of different contexts, three specific to biological networks and two more general algorithms. We found substantial sensitivity of clustering results to the injection of small amounts of noise into the networks. We then reasoned that individual clusters that are robust to small perturbations are more likely to represent biologically or sociologically coherent communities. To this end, we developed the tool *clust.perturb*, an R package (https://github.com/GregStacey/clust-perturb) and Shiny web application (https://rstacey.shinyapps.io/clust-perturb-tool/), that measures the reproducibility of clusters, and nodes within clusters, over multiple iterations of network perturbation. We show that *clust.perturb* can accurately predict which clusters are likely to be reproduced in real-world situations where networks vary (experiment-to-experiment changes), motivating its use as an additional computational step when constructing clusters from networks.

## METHODS

### Datasets

We analyzed the robustness of several clustering algorithms, using both undirected graphs and raw proteomic data as inputs. First, we perturbed networks by randomly removing and adding edges. To provide a broad range of networks, we clustered two biological networks and a social network (Supp. Fig. 1):

1. A protein-protein interaction network (“interactome”), composed of a list of literature curated protein-protein interactions (CORUM, https://mips.helmholtz-muenchen.de/corum/, downloaded September 2018) (Giurgiu et al. 2019). Because CORUM is published as a list of protein complexes, not pairwise interactions, we first reduced the 2824 CORUM complexes among human proteins to their pairwise network. This produced a network of 39563 protein-protein interactions (edges) between 3645 unique proteins (nodes). The original protein complexes were used as a ground-truth cluster set.
2. A drug-drug interaction network, which is a subset of the DrugBank database that lists drug pairs with known interactions, i.e. drug pairs that have unwanted side effects when taken together (DrugBank, https://snap.stanford.edu/biodata/datasets/10001/10001-ChCh-Miner.html, downloaded May 2019) (Wishart et al. 2018). This network consists of 48514 drug-drug interactions (edges) between 1514 unique drugs (nodes).
3. A social network, consisting of anonymized emails between members of a European research institution (email-Eu, https://snap.stanford.edu/data/email-Eu-core.html, downloaded May 2019) (Yin et al. 2017; Leskovec, Kleinberg, and Faloutsos 2007). Nodes in this network represent members of the institution, and are connected by an edge if either person sent the other at least one email (16063 edges between 1005 nodes). The original network was directed and contained self-interactions. For the purposes of this study we removed self-interactions and modified it to be undirected. Each research institute member was a member of exactly one of 42 departments, and these department affiliations were used as a ground-truth cluster set.

All three networks were unweighted (i.e. all edge weights=1), undirected, and had no self-interactions. The CORUM and email-Eu networks had ground-truth cluster assignments.

Second, in order to specifically evaluate the impact of experimental noise on cluster assignment from biological data, we also clustered four co-fractionation datasets collected by our laboratory (Scott et al. 2017; Kristensen, Gsponer, and Foster 2012; Scott et al. 2015; Kerr et al., 2019). These datasets were collected to reconstruct protein-protein interaction networks, using co-fractionation as evidence of protein interaction. They were collected across different species and experimental conditions and represent a broad range of co-fractionation experiments: Kristensen et al. (2012) studied the response of HeLa cells to epidemial growth factor over three biological replicates; Scott et al. (2015) studied HeLa cell response to Salmonella infection, four replicates; Scott et al. (2017) studied apoptotic Jerkat cells, three replicates; and Kerr et al. (2019) studied Hela cell response to interferon stimulation, four replicates. Each dataset is composed of thousands of co-fractionation profiles collected under two experimental conditions and repeated in at least three replicates. In this study we treated each combination of replicate and experimental condition separately, of which there were 28 in total. To generate an interactome network from each replicate/condition, we used the PrInCE analysis pipeline with default settings (Stacey et al. 2017). As in previous work (Skinnider et al. 2018, Stacey et al. 2018), our intent in so doing is not necessarily to argue that PrInCE represents the single most accurate approach for protein-protein interaction inference from co-fractionation data, but rather that it is sufficiently representative of the kinds of machine-learning approaches used within the field (e.g., Havugimana et al. 2012, Wan et al. 2015, Hu et al. 2019) that our conclusions will generalize more broadly. Three replicate/condition datasets produced interactomes with fewer than 50 pairwise interactions, likely because of poor data quality, and these were not analyzed in this study. Before adding noise, the remaining 25 datasets used for this study produced interactomes with between 143 and 14358 interactions (6139 +/- 3989, mean +/- st.d.).

### Experimental methodology

In order to test the effects of network noise on clustering results, we added network-level noise by randomly rewiring network edges, and clustered both noised and original networks using graph-based clustering algorithms. That is, a certain number of edges were removed from the network and replaced with the same number of edges not previously contained in the network. Thus, the size of the network was preserved while the fraction of rewired edges was varied. The fraction of rewired edges was calculated as a false positive rate, which is equal to the number of rewired edges divided by the total number of edges in the network. In order to avoid confusion with a network’s inherent false positive rate, which comes from mistakes in the original edge list, we use the term “network noise level” to refer to the false positive rate of *in silico* changes made to the network. To track the effects of network noise on clustering, we compared clustering results from noised networks to the clustering results from the original network (Fig. 1A). We repeated this noised-to-original comparison for each clustering algorithm.

**Figure 1.**
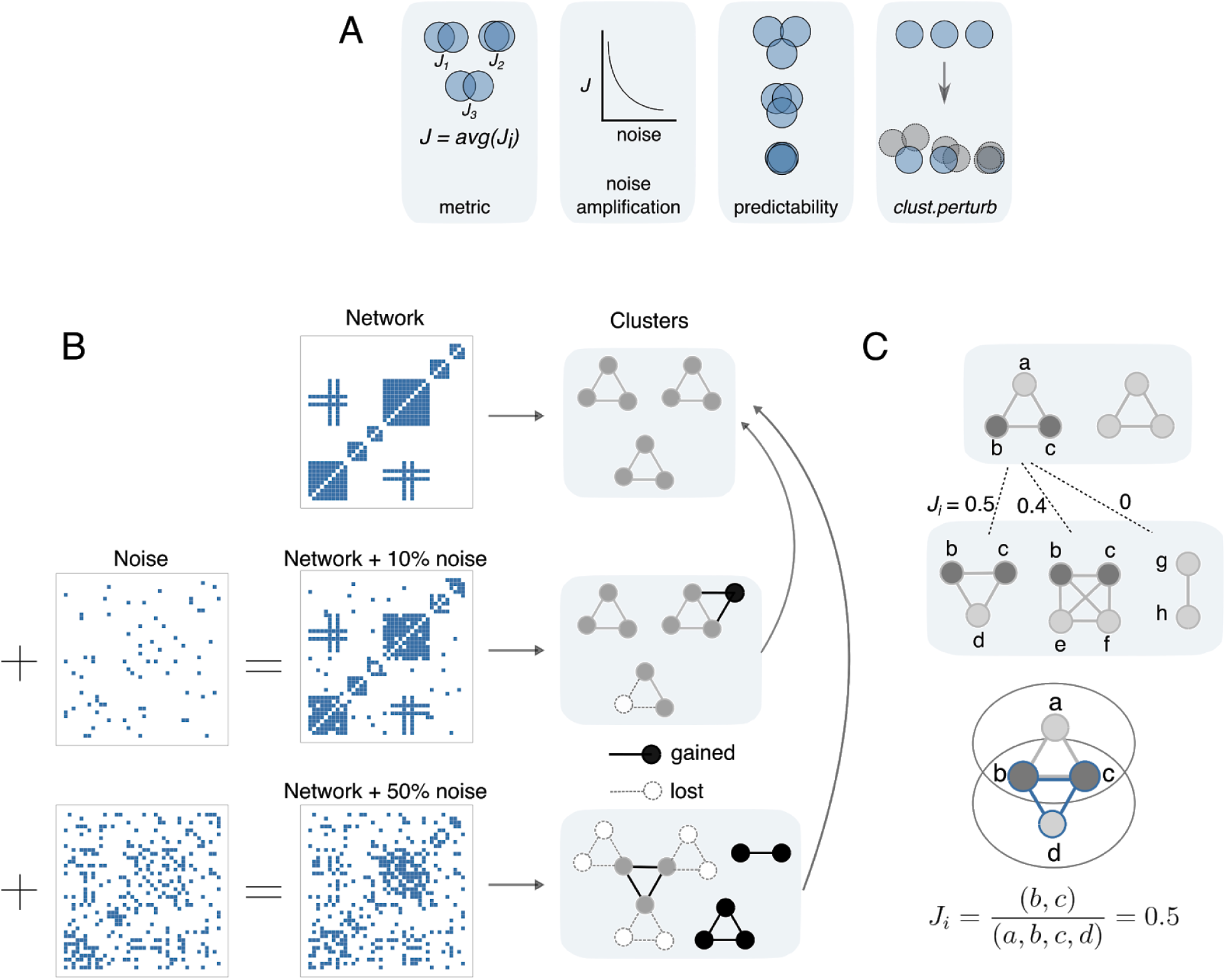
Testing the effect of network noise on clustering. A) Primary findings of this study. B) Overview of methodological design. Networks with varying amounts of noise are clustered and results are compared to clusters derived from the original networks. C) Cluster-wise comparisons are made using the Jaccard index.

In addition to rewiring, we ran limited analyses with different levels of added or removed edges to examine whether our conclusions are specific to a particular type of noise (that is, false positives or false negatives). In these cases, a fraction of edges were removed, or edges not originally in the network were added, or both in differing amounts. Similar to rewiring, the number of added or removed edges was proportional to the size of the network. Noise levels analyzed were *fnoise =* 0%, 1%, 2%, 5%, 10%, 15%, 25%, 50%, and 100%. Since this produces 81 noise combinations, we also calculated the overall level of injected network noise as Δ*edges* = *fadd + fremove*, i.e. the sum of the added and removed fractions.

The effects of experimental noise on clustering results were measured in much the same way, except that noise was added to co-fractionation profiles prior to generating a network via PrInCE, rather than to the network directly. Noise was added to co-fractionation profiles by adding a normally distributed random number to the log-transformed value of each data point, with standard deviation equal to the noise level. That is, for each data point,

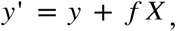

where *y*’ is the noised log-transformed co-fractionation data, *y* is the original log-transformed co-fractionation data, *f* is the co-fractionation noise level, and *X* is a normally distributed random number with mean zero. Log-transformation was applied to ensure positive values, and because co-fractionation values tend to follow an approximately log-normal distribution.

### Choosing a cluster-wise and set-wise similarity metric

Many different metrics have been proposed to measure the similarity of two sets of clusters (Gates et al. 2017). These metrics implicitly trade off rewarding intra-cluster edges (true positives) and penalizing inter-cluster edges (false positives). Previous work has shown that this tradeoff leads different cluster similarity metrics to measure distinct aspects of cluster similarity, and to display non-intuitive behaviour (Gates et al. 2017). Therefore, we examined which set-wise similarity metrics, if any, adhered to intuitive notions of “cluster set similarity”, such as measuring complete similarity when sets are identical and less than complete similarity when they are not. Set-wise metrics assign a similarity value to entire sets of clustering results, i.e. a single value for all clusters. This in contrast to cluster-wise metrics, which assign a similarity score to each cluster. Set-wise metrics are often computed as an average of cluster-wise metrics.

We analyzed whether set-wise metrics matched intuition in four scenarios:

1. A similarity score of 1 is assigned to identical cluster sets and decreases as the number of non-identical cluster assignments increases.
2. The score is not biased by the number of clusters in either set.
3. The score is not negatively affected by nodes participating in multiple clusters (“moonlighting”).
4. The score penalizes situations where sets are non-identical because of missing nodes in one set.

These were tested using simulated clustering sets of 1000 nodes assigned to 100 equal-sized clusters, prior to manipulation in each condition. Our analysis parallels that of (Gates et al. 2017). In this simulation we tested six commonly-used set-wise cluster similarity metrics:

- Normalized mutual information (NMI) (McDaid, Greene, and Hurley 2011)
- Adjusted rand index (ARI) (Hubert and Arabie 1985)
- Geometric accuracy (GA) (Brohée and van Helden 2006; Freytag et al. 2018; Nepusz, Yu, and Paccanaro 2012)
- Maximum matching ratio (MMR) (Nepusz, Yu, and Paccanaro 2012)
- F-measure
- Jaccard index

### Maximum Jaccard index (*J*) and simple counting statistics

To measure whether cluster *i* in cluster set 1 is also contained in cluster set 2, we quantify cluster-wise similarity using the maximum Jaccard index, which is the number of nodes in common between two clusters divided by the total number of unique nodes in the two clusters (Fig. 1B). The cluster-wise Jaccard index *Ji* is calculated as

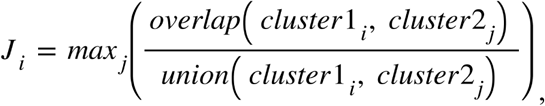

where *cluster1* is the noised cluster set, *cluster2* is the original cluster set, and *cluster1*_*i*_ and *cluster2*□ are single clusters from those sets. That is, the similarity score *Ji* of a cluster from a noised cluster set is equal to the maximum Jaccard index between that cluster and any cluster in the original set. Set-wise similarity *J* is quantified by averaging *Ji* over all clusters in the noised cluster set. That is,

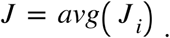

In addition to metric *J*, we also employed simple counting statistics to measure the difference between cluster sets. These include the number of gained nodes (nodes present in a noised cluster that are not present in the best-match original cluster), the number of lost nodes (nodes not present in a noised cluster which are present in the best-match original cluster), and the number of rearranged cluster edges. The latter is defined as the absolute number edges different between a noised and original cluster set, i.e. the number of edges one would need to add to or remove from the original cluster set to make it identical to the noised set.

### Clustering algorithms

We analyzed the results of five clustering algorithms: Markov clustering (“MCL”) (van Dongen 2000), ClusterONE (“CO”, java implementation at www.paccanarolab.com) (Nepusz, Yu, and Paccanaro 2012), a two-stage clustering algorithm of CO followed by MCL (“CO+MCL”), k-medoids (“k-medoids”, R function *pam*), and walktrap clustering (“walktrap”, R function *walktrap.community*) (Pons and Latapy 2005). We chose these algorithms to provide a broad range of algorithms, including those popular for clustering interactomes (CO, MCL, CO+MCL) as well as more general clustering algorithms (k-medoids, walktrap). CO+MCL has been used specifically to avoid large clusters sometimes predicted by CO (Wan et al. 2015; Drew et al. 2017). The k-medoids algorithm is similar to k-means. k-medoids was used instead of k-means, because the R package *pam* allows analysis to start from an adjacency matrix, whereas common R implementations of k-means do not. This is important since our method adds noise to adjacency matrices (Fig. 1A).

Parameters for all clustering algorithms were chosen by grid search optimization. Parameter optimization was performed with the original networks. Since CORUM and email-Eu had ground truth cluster assignments, we chose the parameter sets that maximized set-wise *J* between clusters derived from the original network and the ground truth clusters. For DrugBank, we chose the parameter set that maximized the silhouette score of the cluster assignments, which selects well-separated clusters whose members are tightly connected by pairwise edges. Parameter ranges and optimized values are given in Table 1. Parameters are:

- P (CO, CO+MCL): Penalty term modeling the number of unknown edges in a network.
- Dens (CO, CO+MCL): Minimum cluster density (fraction of filled edges between cluster members).
- I (CO+MCL, MCL): Expansion parameter.
- Nclusters (k-Medoids): Explicitly controls number of clusters.
- Steps (walktrap): Number of random walks.

**Table 1.** Parameter choices and ranges for clustering algorithms.

| Clustering algorithm | Optimal parameter choices (CORUM) | Parameter ranges | Non-optimal parameter choices |
| --- | --- | --- | --- |
| CO | P = 500<br>Dens = 0.1 | P: [1, 50, 100, 500, 5000]<br>Dens: [0, 0.1, 0.2, 0.3, 0.4] | P = 2<br>Dens = 0.3 |
| CO+MCL | P = 500<br>Dens = 0.1<br>I = 4 | P: [1, 50, 100, 500, 5000]<br>Dens: [0, 0.1, 0.2, 0.3, 0.4]<br>I: [1, 2, 4, 8, 15, 20, 50] | P = 2<br>Dens = 0.3<br>I = 2 |
| MCL | I = 2 | I: [1, 2, 4, 8, 15, 20, 50] | I = 4 |
| k-Med | Nclusters = 1500 | Nclusters: [50, 100, 250, 500, 1000, 1500, 2000, 5000] | Nclusters = 1000 |
| walktrap | Steps = 2 | Steps: [2, 4, 6, 8, 10, 12] | Steps = 10 |

For full parameter explanations see (Nepusz, Yu, and Paccanaro 2012) (CO), (van Dongen 2000) (MCL), (Jin and Han 2016) (k-Medoids), and (Pons and Latapy 2005) (walktrap).

### *clust.perturb*: a tool for assessing cluster reliability

*clust.perturb* is both an open source R package (https://github.com/GregStacey/clust-perturb) and web application (https://rstacey.shinyapps.io/clust-perturb-tool/, R shiny) designed to assess cluster reproducibility by detecting the tendency for clusters to change after random perturbations are applied to the clustered network. It is designed as a general-purpose wrapper to clustering algorithms, which will return both the original clusters and measures of the reproducibility with which individual clusters and proteins are detected by the algorithm. The R package can be used to assess clusters from any clustering algorithm, while the web application uses three default clustering algorithms (hierarchical, MCL, and k-Medoids). Networks are perturbed by rewiring edges as described in this paper, and the networks are input as edge lists. *Clust.perturb* takes three input parameters and returns two outputs. The input parameters are: clustering algorithm; number of iterations; and noise level, quantified as the fraction of network edges that are rewired. The outputs are: *repJ*, a measure of cluster reliability, which is equal to the cluster’s average *Ji* over all noise iterations; and *fnode*, a measure of node reproducibility within a cluster, which is equal to the frequency with which that node occurs in best-matched clusters divided by the number of noise iterations.

## RESULTS

In this study we investigated the degree to which graph-based clustering of biological and social networks are contaminated by network-level noise. To do so, we sought to address four questions (Fig. 1A). First, we established a suitable metric to measure changes in clustering solutions after injection of noise into a network. Second, we used this metric to demonstrate that clustering amplifies network noise, i.e. the ratio of network level noise to set-wise Jaccard index *J* is greater than 1, such that injection of a small degree of noise into a network can result in dramatic changes to its clustering. Third, we demonstrated that it is possible to predict which clusters will be most affected by noise. Finally, we developed a tool (*clust.perturb*) that estimates the reproducibility of cluster assignments.

### An intuitive metric for measuring clustering similarity

Measuring the effects of noise on clustering results requires a metric for quantifying the difference between two cluster sets. Because quantifying clustering similarity requires choosing how to reward true positives (within-cluster edges) and penalize false negatives (between-cluster edges), this task can be surprisingly ambiguous. There exist many commonly used cluster similarity metrics (Xu and Tian 2015) with biases that result in unintuitive behaviour (Gates et al. 2017). Therefore, we tested a number of commonly used metrics with the goal of ensuring that we are indeed measuring an intuitive notion of “cluster set similarity”. Confirming previous results (Gates et al. 2017), we found that most metrics failed to adhere to intuition (Fig. 2). For example, geometric accuracy (GA) and normalized mutual information (NMI) can both measure disagreement between completely identical sets (in cases of “moonlighting”, i.e. where nodes are assigned to multiple clusters, Fig. 2C) and measure perfect agreement between non-identical sets (when a cluster set contains nodes not contained in the other set, Fig. 2D). However, the maximum Jaccard index *J* (Fig. 1B) behaved intuitively in all situations tested. This measure also has the benefit of measuring both cluster-wise and set-wise similarities (*J* and *Ji* respectively). We therefore use *J* and *Ji* throughout this study.

**Figure 2.**
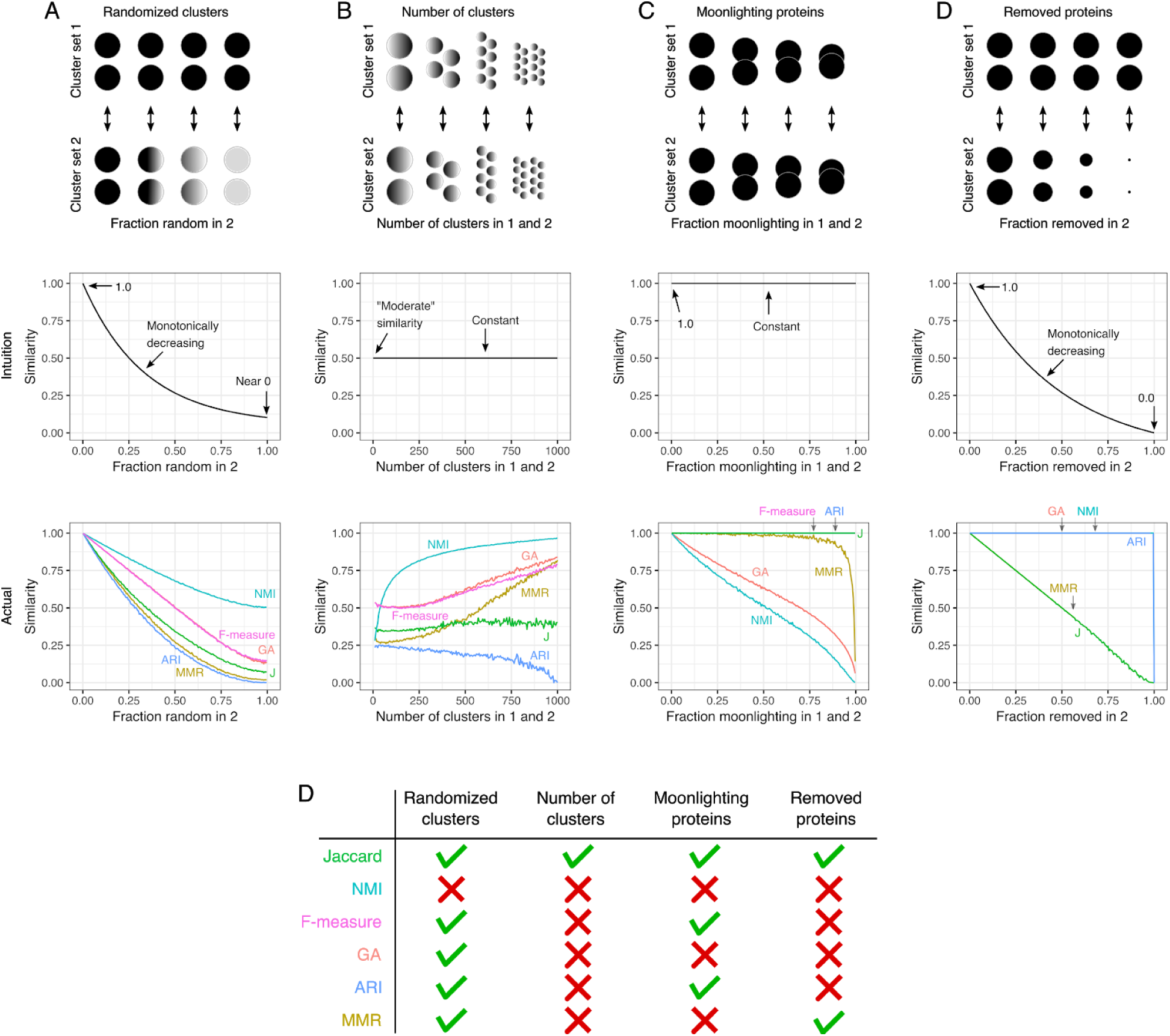
Maximum Jaccard index (*J*) is consistent with intuitive notions of “cluster set similarity” in all four cases. In all cases 1000 nodes are assigned to clusters, except D where the number of nodes in set 2 is varied. A) Set 1: 1000 nodes assigned to 100 clusters of equal size. Set 2: identical to Set 1, aside from a variable fraction of nodes randomly assigned to an existing cluster. B) In all comparisons 50% of cluster assignments are identical, and the number of clusters is varied. Set 1: 500 nodes assigned to a variable number of clusters of equal size, with the remaining 500 nodes randomly assigned to an existing cluster. Set 2: identical to Set 1 for the first 500 nodes, but different random assignments for the remaining 500 nodes. C) Cluster Sets 1 and 2 are identical in all comparisons, and the number of nodes assigned to multiple clusters is varied to simulate “moonlighting” nodes. Sets 1 and 2: 1000 potentially non-unique nodes are assigned to 100 clusters of equal size. At fraction=0 all nodes are unique, while at fraction=1 all nodes are the same node. D) Set 1: 1000 nodes nodes assigned to 100 clusters of equal size. Set 2 is identical to Set 1 before a fraction of Set 2 nodes are removed. At fraction=0 both sets are identical, while at fraction=1 Set 2 is empty.

### Clustering amplifies network-level noise

Since graph-based clusters are generated from a network, one would expect that changes to the network would also lead to changes in clusters. Indeed this is the case: as network-level errors increase in the CORUM network, cluster sets both lose and gain proteins when compared to sets of clusters derived from the error-free network (Fig. 3A). This response to injected network noise was quantified by our chosen cluster-wise and set-wise metrics (*Ji* and *J*, respectively) (Fig. 3B). The injected network errors in this case are random connections between proteins that do not necessarily share any biological role, and the removal of true positives, which should be accompanied by a loss of biological plausibility in the clustered proteins. Therefore as a control analysis, we confirmed that clusters derived from noised networks are less enriched for Gene Ontology (GO) terms than clusters derived from the CORUM network without added noise (Supp. Fig. 2).

**Figure 3.**
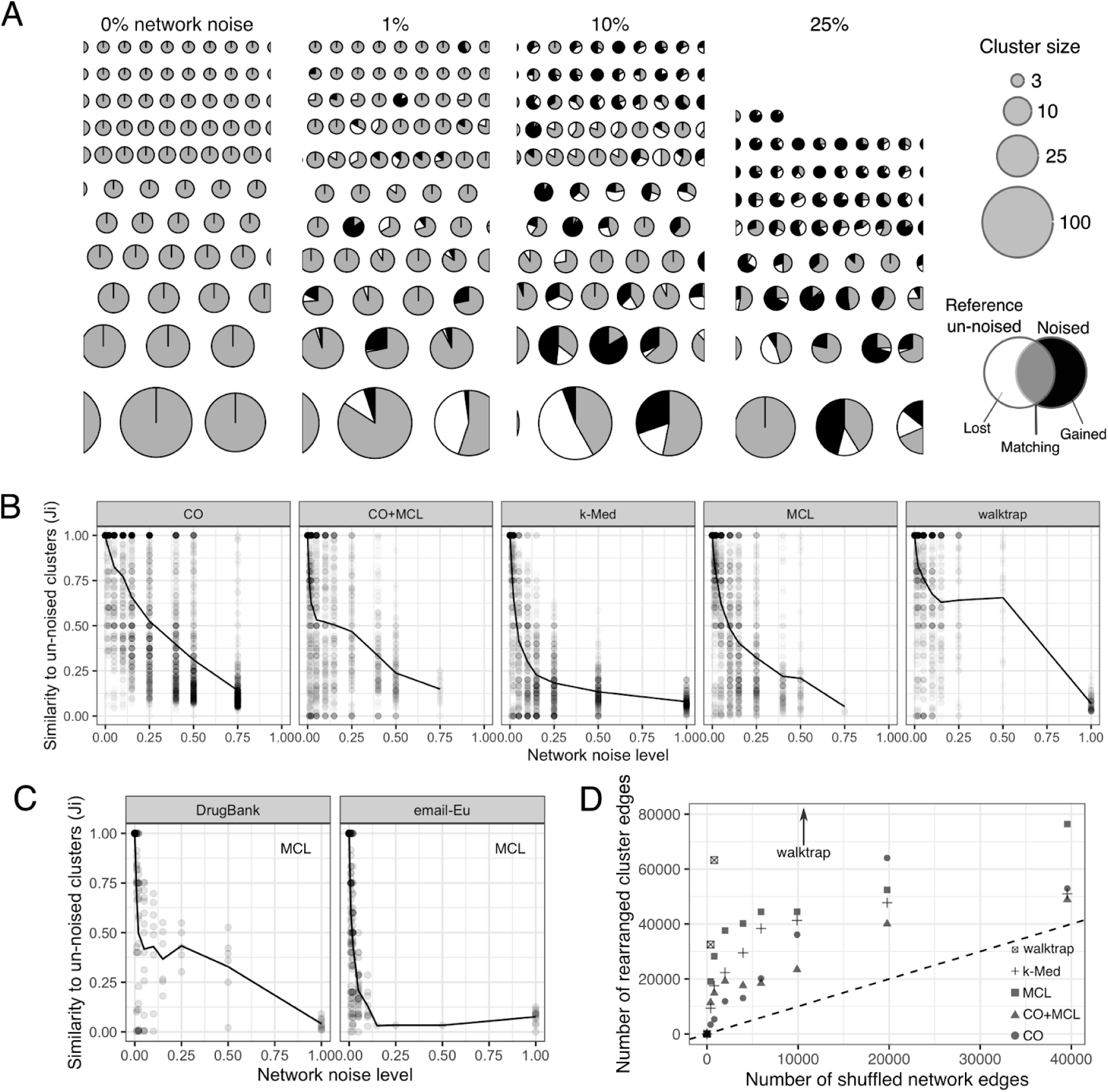
Clustering amplifies network noise. A) Subset of the cluster sets produced by clustering the CORUM network with 0-25% network noise. Colour shows the fraction of proteins overlapping with original clusters, proteins lost after adding noise, and proteins gained. Clusters were made with the MCL algorithm. B) Quantifying the effects of CORUM network noise using set-wise metric *J* (line) and cluster-wise metric *Ji* (scatter) Line shows average *Ji* (*J*). C) Quantifying the effects of network noise on DrugBank and email-Eu networks. Algorithm MCL. D) Comparing the number of altered network edges to the number of rearranged cluster edges.

While it is to be expected that large amounts of injected network noise lead to changes in clustering results, we find that this is the case for even small amounts of injected network noise. For all clustering algorithms, an alteration of 1% of the binary interactions (edges) in the CORUM network resulted in the alteration of at least 13% of clusters in a set (53/406 clusters, CO) and at most 58% of clusters (182/314 clusters, CO+MCL). Measuring the set-wise change, an introduction of 1% network noise lead to a 4-29% change in clustering results as quantified by *J* (*J*=0.96 to *J*=0.71); and a 2% noise resulted in set-wise clustering differences between 8-41% (Fig. 3B). Additionally, there was a positive relationship between cluster size and reproducibility, as shown by multiple linear regression between *Ji* (dependent variable) and network noise level, algorithm, and cluster size (beta=0.2 10-4, p=1.3 10-6; model R2=0.42). That is, small clusters were significantly less reproducible. Importantly, the variability of cluster sets measured here is due to network noise and not inherent randomness in the clustering algorithms: all of the algorithms analyzed here are deterministic, meaning the clustering does not vary if the network is unchanged (see Figure 3B, Ji=1 when noise=0).

Amplification of network noise by clustering was also seen when clustering the DrugBank and email-Eu networks (Fig. 3C and Supp. Fig. 3). Similarly to clustering results from the CORUM network, small network changes (rewiring 1-2% of the edges) lead to substantial changes in clustering results, including total loss of some clusters. This is reflected in a similar level of noise amplification to CORUM clustering, as 1% network noise reduced *J* by 32-57% (DrugBank) and 10-40% (email-Eu). Therefore, we conclude that sensitivity to noise is a robust property of graph-based clustering, as applied to many different types of networks.

Clustering appears to be sensitive to network noise when measured by *J*. However, set-wise metrics can be misleading (Fig. 2), so we also sought to confirm that substantial clustering rearrangements were occurring through visual inspection. Supplementary Figure 4A-E shows visually the results of clustering the binarized CORUM network via MCL and the extent of gained/lost proteins at various noise levels. Consistent with conclusions based on metric *J*, after 1% of the binary interactions were rewired, more than a third of clusters underwent some rearrangement (102/305 clusters), with all clusters losing 0.88 proteins and gaining 1.82 proteins on average (Supp. Fig. 4F). We also counted the number of lost and gained cluster edges, i.e. the number of edges one would need to alter in the original cluster set in order to arrive at a noised cluster set (Fig 3F). This permitted us to directly compare the number of rewired network edges to the number of rearranged cluster edges. We saw that at low levels of network noise (1% network noise level), rewiring a single network edge produced 38.2 rearranged cluster edges on average, consistent with clustering amplifying network noise.

Network noise commonly involves both the absence of truly occurring edges (false negatives) and the presence of spurious edges (false positives). In the rewiring experiments above, we simulated the addition of both false positives and false negatives simultaneously. We also asked whether the consequences of network noise for clustering robustness varied when false positives and false negatives were added in varying proportions. We saw that regardless of whether edges were removed, added, or rewired, clustering consistently amplified low levels of injected network noise (Supp. Fig. 5B, red vs. blue). However, at higher noise levels, we found that edge removal generally had a greater effect than edge addition.

Finally, we considered the possibility that the sensitivity to network noise that we observed could be specific to the set of clustering algorithm parameters that yielded the optimal clustering solution. Although it is unlikely that this result is unique to a parameter set, given that we employed five parameter sets that were chosen by two optimization schemes (optimal ground truth similarity and optimal silhouette score), we wanted to test whether noise sensitivity persisted when clustering the same network with the same algorithm but specifically selecting a different, non-optimal parameter set (Table 1). With non-optimal parameters, 1% network noise lead to a minimal rearrangement of *J*=0.85 (CO) and a maximal rearrangement of *J*=0.66 (CO+MCL), comparable to values *J*=0.95 and *J*=0.66 for cluster sets using optimized parameters (Fig. 3). Therefore, sensitivity to network noise appeared to be a general feature of the clustering algorithms studied here, rather than a consequence of parameter optimization.

### Clustering results derived from experimental datasets are also poorly reproducible

The binarized CORUM network may not be representative of a typical experimentally-derived network, for a number of reasons. For instance, CORUM is relatively complete compared to an experimental dataset, with fewer false negatives (Stacey et al., 2018), and many (tens of thousands) of binary interactions. Therefore we sought to confirm our findings using clustering results derived from experimental co-fractionation datasets, which typically have fewer interactions that are more sparsely connected. In this analysis, we added noise directly to the underlying proteomic data rather than the network (Fig. 4A), then subsequently performed network inference on the noisy proteomic data, in order to investigate the impact of experimental noise on clustering. Consistent with previous results from the binarized CORUM network, injecting relatively insignificant levels of noise into experimental co-fractionation datasets resulted in substantially altered clustering results (Fig. 4B). For example, although co-fractionation profiles with 1% added noise were highly similar to profiles without added noise (average Pearson R^2^ = 0.9996, Fig. 4D), clustering these two noise levels could produce clustering sets with large differences (Fig. 4B, CO+MCL *J*=0.68), the magnitude of which were comparable to those observed in CORUM. Importantly, injecting small amounts of noise also had small effects on the predicted interactome network, as interactomes derived from co-fractionation profiles with 1% added noise were largely similar to interactomes derived from profiles with no noise (Jaccard = 0.96, Fig. 4D), and similarly for 2% added noise (Jaccard = 0.93). That is, slight alterations to experimental data could abolish a large proportion of clusters, while having a lesser impact on the interactomes from which the clusters are derived. This is consistent with the results obtained from analysis of CORUM, and reflects the amplification of both network-level noise, as well as noise in the underlying experimental data, by graph-based clustering.

**Figure 4.**
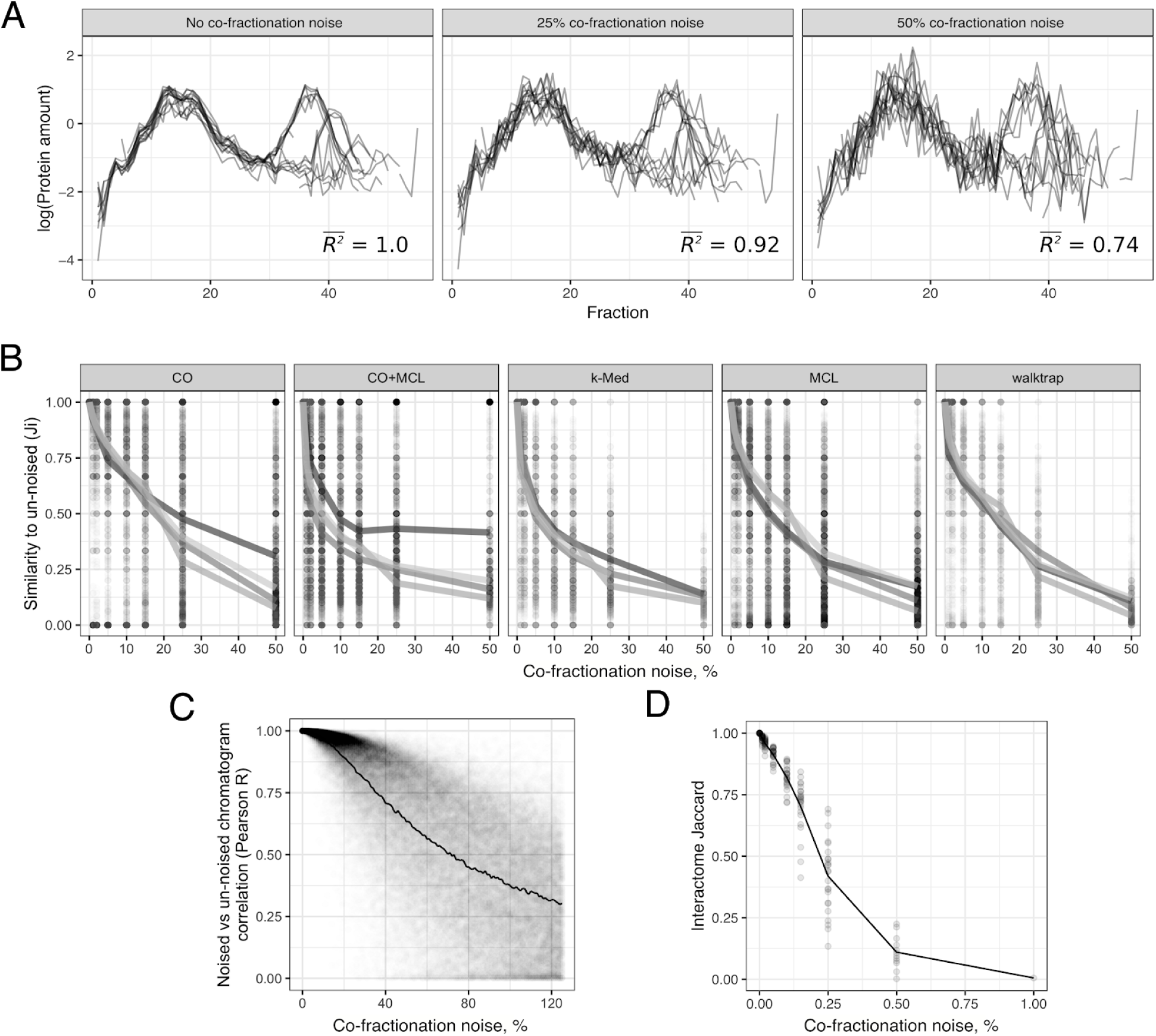
Noise in experimental datasets also affects clustering results. A) Co-fractionation profiles of 26S proteasomal proteins with no added noise (left), 25% noise (middle), and 50% noise (right). Average R^2^ values shown are calculated between each co-fractionation profile before and after adding noise. In the “No noise” case these are the same profiles, hence perfect correlation. B) Effects of adding co-fractionation noise on clustering results, measured with *J* (lines) and *Ji* (points). C) Quantifying the degree to which added co-fractionation noise degrades co-fractionation profiles, as measured by Pearson correlation between noised and original co-fractionation profiles. D) Effect of co-fractionation noise on interactomes, as measured by Jaccard index between noised and original interactome. Each dot is a dataset, and the line shows the average.

### The response of individual clusters to network noise is reproducible

Our results thus far indicate that the clustering algorithms studied here are globally sensitive to small levels of noise in the underlying networks. However, not all clusters are equally affected by this sensitivity to noise. For example, adding 5% noise to the CORUM network and clustering with k-Medoids produced a cluster set with moderate similarity to the original set (J = 0.42 at 5% noise, Fig. 2B, k-Med panel). However, within that set, some clusters remained unchanged (*Ji* = 1, top) while others were entirely removed (Ji = 0, bottom). We investigated the consistency of this pattern, that is, whether some clusters tended to be more stable in response to network noise than others. We reasoned that, if some clusters are consistently reproducible in response to simulated network noise, then it may be possible to identify clusters that will be robust to future, real-world alterations of the network, e.g. the collection of data in future experiments.

To quantify cluster stability, we performed multiple iterations of noise injection and clustering using the CORUM network, and then calculated *Ji* between the original cluster set and each iteration. That is, for each of the original clusters we calculated *N* values of *Ji*, where *N* is the number of noise iterations. If some clusters are consistently stable (or unstable) in response to noise, the *Ji* values should be consistently high (or low) across noise iterations. This is indeed what we observe. Measuring this consistency as a correlation in the Jaccard index between iterations, Figure 5A shows *Ji* values for two iterations of clustering, using the ClusterONE algorithm, across independent noise injections. These iterations are significantly correlated (R = 0.72, p<10-15, Pearson correlation; ClusterONE). For all clustering algorithms studied here, the reproducibility of individual clusters was correlated between random noise iterations, with ClusterONE having the highest correlation and MCL the lowest (Fig 5B). Across algorithms, a cluster’s reproducibility correlates with its density, i.e. fraction of intra-cluster edges (Spearman R = 0.35, p<10-16). Taken together, this suggests that a cluster’s tendency to “break” in response to injection of network noise is predictable, and that more reproducible clusters are more supported by intra-cluster edges in the underlying network.

**Figure 5.**
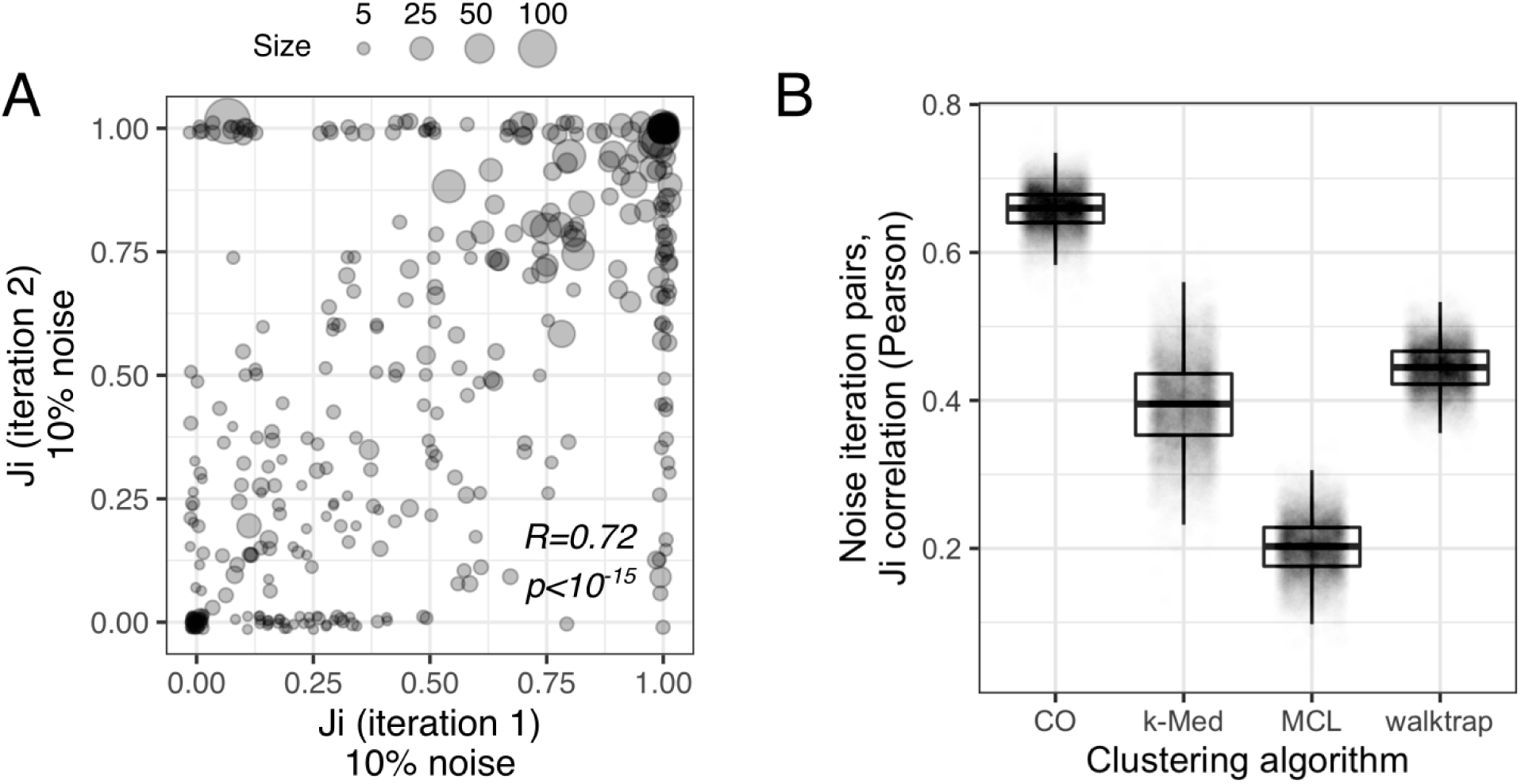
Response of individual clusters to network noise is consistent across noise iterations. A) *Ji* values between clusters of the CORUM network (ClusterONE) and two noise iterations (ClusterONE plus 10% shuffling). Pearson correlation. B) Pearson correlation values between pairs of noise iterations for four clustering algorithms.

### A software tool for predicting cluster reproducibility (*clust.perturb*)

Some clusters are consistently robust when the network is randomly altered *in silico*. If the simulated network noise is representative of real-world network alterations, it should be possible to predict the effect of future alterations to the network, and thereby identify robust clusters. To this end we developed *clust.perturb*, an R-based tool for measuring cluster reproducibility by randomly perturbing the network. *clust.perturb* takes a network (.*csv* or .*tsv* edge list) as input and returns two scores, *repJ* and *fnode*, which quantify the reproducibility of clusters and the reproducibility nodes within clusters, respectively (Fig. 6A). Following the previous analysis, *clust.perturb* first clusters the network, then performs *N* iterations of clustering with network noise, yielding *N* values of *Ji* for each cluster from the original cluster set. *repJ* is then calculated as the average of these *Ji* values. For example, ClusterONE identifies a thirteen-protein cluster in the CORUM network loosely corresponding to the G alpha-13-Hax-1-cortactin-Rac complex (Fig. 6B, left). Over multiple network noise iterations (Fig. 6B) seven proteins in the original complex tend to remain co-clustered (Fig. 6C, top yellow) while the other six proteins do so less consistently. On aggregate this cluster is partially reproduced, reflected by its reproducibility score of *repJ* = 0.61 (100 iterations). Other clusters are effectively “lost” when the network is altered (Fig. 6D, repJ=0.41), whereas others still remain largely unchanged (Fig. 6E, repJ = 0.87).

**Figure 6.**
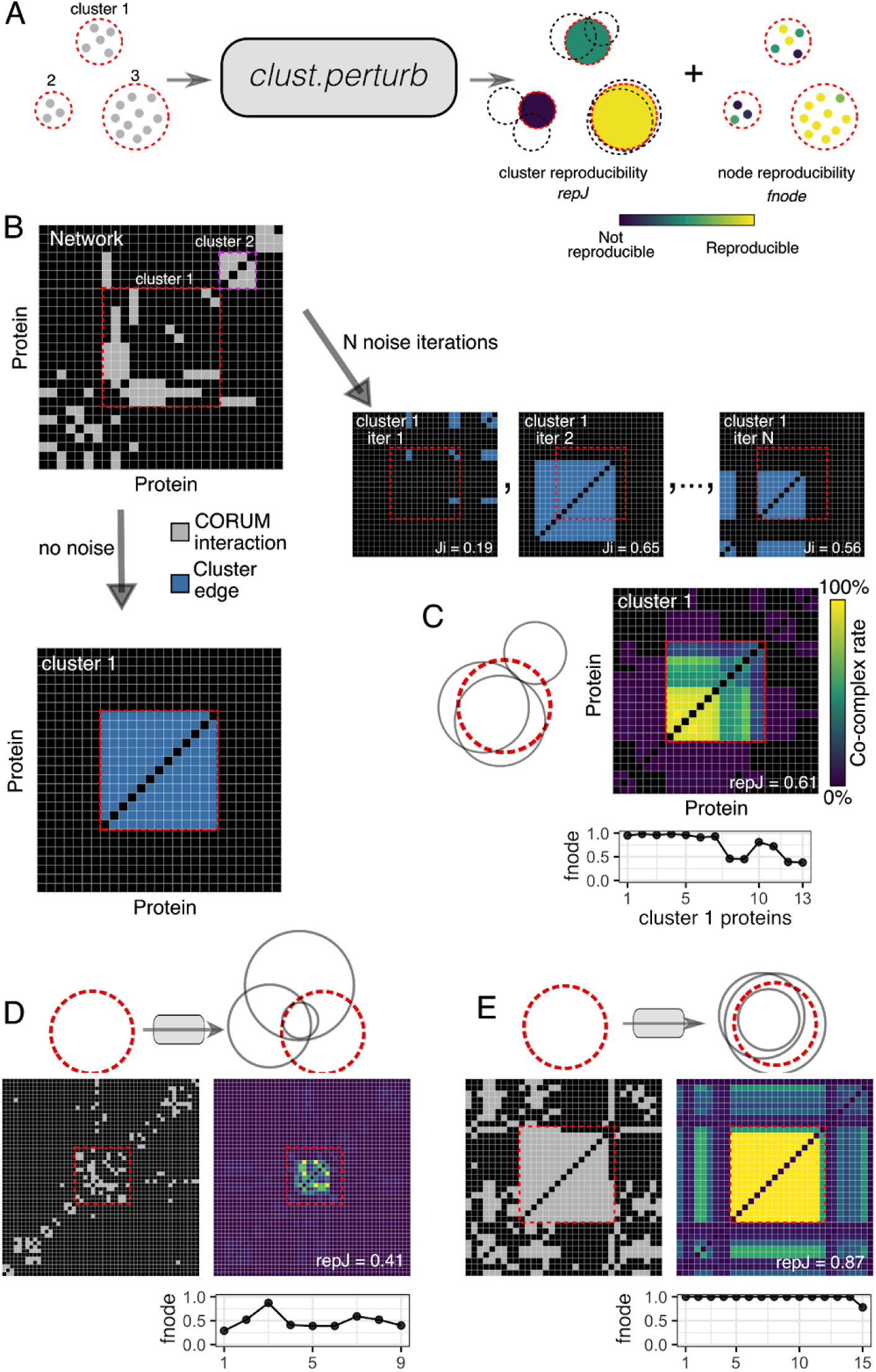
*clust.perturb* measures cluster and node reproducibility. A) Schematic. *clust.perturb* clusters a network with and without noise. A cluster’s reproducibility is its average overlap with best-match clusters across noise iterations (*repJ*). Node reproducibility is the frequency with which a cluster’s nodes appear in best-match noise clusters, divided by the number of noise iterations (*fnode*). B) Network adjacency matrix showing a subset of the CORUM network (grey) and a 13-protein cluster identified in the network (cluster 1, blue). Adjacency matrices for best-match clusters from 3 noise iterations are also shown (10% noise). C) Average adjacency matrix for cluster 1 across 100 noise iterations. *fnode* values for each protein in cluster 1 are shown. Venn diagram shows overlap between cluster 1 (red) and best-match clusters from three noise iterations (grey). D) Cluster with low reproducibility. E) Cluster with high reproducibility. ClusterONE, 10% network noise.

In addition to scoring the reproducibility of each cluster, *clust.perturb* also scores the reproducibility with which each node (e.g. protein) within a cluster is associated with that cluster, by counting the frequency with which it is re-clustered across noise iterations. A score is assigned to each node (*fnode*) based on the frequency with which it occurs in the closest matching noised cluster. For example in Figure 6B, *fnode* values close to 1 reflect the fact that these proteins are present in the best-matching cluster in nearly all noise iterations, while lower *fnode* values identify proteins that “drop out” of clusters.

*clust.perturb* requires two parameters in addition to the network and choice of clustering algorithm: number of noise iterations (*N*) and magnitude of network noise (*m*). We sought to establish sensible defaults for these parameters. Because the running time scales linearly with *N, clust.perturb* will be time-intensive if the original clustering algorithm is time-intensive. To find the minimum iterations necessary to adequately estimate a cluster’s reproducibility, we took *N=*100 iterations as a “final” value, and we calculated how quickly *repJ* converged to that value over successive iterations. After a single iteration the average absolute error for a given *repJ* was 0.16, and after 25 iterations it was 0.03 (ClusterONE, 10% network noise) (Fig. 6A). For all algorithms a single iteration was sufficient to estimate the final *repJ* with a median error of 0.15 (Fig. 6B). Thus, we find that *repJ* converges relatively quickly, and a few iterations are often adequate to accurately approximate cluster robustness. Next, the noise magnitude *m* should be chosen to properly resolve *repJ* values: if a noise level is too small, most clusters will be unchanged and the resolving power will be poor (left, Fig. 6C), whereas the opposite is true if noise is too great (right). In general, we found a noise level of 10% was sufficient to resolve clusters generated by all algorithms (Fig. 6C centre).

### Validating *clust.perturb* using biological evidence

We next sought to validate the reproducibility measures, *repJ* and *fnode*, calculated by *clust.perturb*. Specifically, do reproducible clusters and nodes correspond to meaningful features of the network? To answer this, we used the original CORUM complexes, which form the ground truth clusters for the network, and Gene Ontology (GO) terms. First, we investigated whether reproducible clusters more closely correspond to ground truth CORUM complexes. To do so we matched each original cluster to its closest COURM complex using maximum Jaccard value. Indeed, for all clustering algorithms we found a significant positive correlation between *repJ* and association with a CORUM complex (Fig. 7A). Similarly, cluster nodes with lower *fnode* scores tended to be proteins outside of the best-match ground truth CORUM complex (Fig. 7B,C). That is, there was a strong, significant association between reproducibility and ground truth for both *repJ* and *fnode* measures.

**Figure 7.**
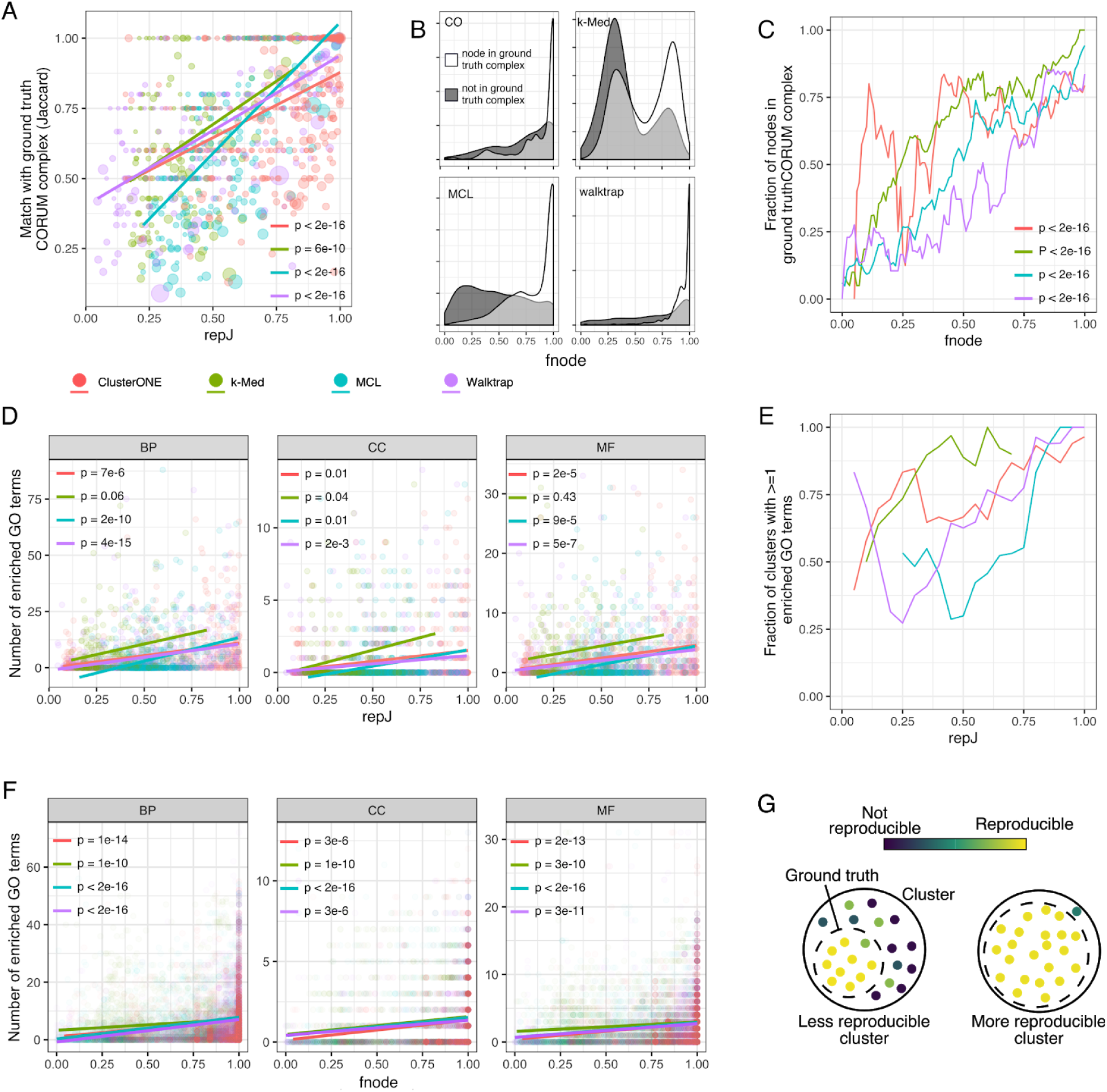
Reproducible clusters and nodes measured by *clust.perturb* are associated with ground truth communities. A) Clusters with high *repJ* values are more closely associated with a CORUM complex, as measured by Jaccard index. P-values are for the *repJ* term in the linear regression model, where *CORUM_jaccard* (dependent) was predicted by *repJ* and *cluster_size* (independent variables). B) Density plots of *fnode* values for nodes in the ground truth CORUM complex (white) and nodes not in ground truth (grey). C) *fnode* values correlate with membership in the ground truth complex. P-values for *fnode* term in logistic regression model where *in_ground_truth* (dependent variable) was predicted by *fnode* and *cluster_size* (independent variables). D) Number of enriched GO terms for a cluster correlates with its *repJ* value. P-values for *repJ* term linear regression model, where *N_enriched_GO* (dependent) was predicted *repJ* and *cluster_size* (independent variables). E) Most clusters with high *repJ* are enriched for at least one GO term. F) Proteins with high *fnode* tend to share GO terms that their assigned cluster is enriched for. G) Nodes “lost” in noise iterations are less likely to be from the ground truth complex, and similarly clusters more tightly associated with a ground truth complex are more reproducible.

We next investigated the association between reproducibility and GO enrichment (hypergeometric test with Benjamini-Hochberg correction, q<0.05; GO terms filtered to >5 and <100 annotations among unique CORUM proteins). For each cluster from the non-noised CORUM network, we counted the number of enriched GO terms in each GO ontology. For all algorithms except k-Med, possibly due to few clusters and therefore small sample size, the number of enriched GO terms was significantly positively associated with a cluster’s *repJ* score, even when controlling for the size of cluster (multiple linear regression, *nGO* dependent variable, *repJ* and *cluster_size* independent; Fig. 7D). Correspondingly, across all algorithms, 94% of complexes with *repJ*>0.8 were enriched for at least one GO term, compared to 50% of complexes with *repJ*<0.5 (Fig. 7E). We next investigated whether a protein’s *fnode* score was associated with GO enrichment by analyzing whether proteins assigned to a cluster shared one of that cluster’s enriched GO terms. Indeed, for all algorithms and ontologies, we found a strong, significant pattern for more reproducible nodes to share the cluster’s enriched GO terms (Fig. 7F). Taken together, these results suggest that reproducible clusters, as measured by *clust.perturb*, more closely align to the network’s ground truth, and that cluster nodes with poor reproducibility are more likely to be spuriously assigned to the cluster (Fig. 7G).

### *clust.perturb* predicts reproducibility across real-world network alterations

If clusters have a consistent response to random, simulated network noise (Fig. 5), then it should be possible to predict which clusters will be robust to future, real-world network alterations. We tested whether this is indeed the case.We constructed two independent networks from each of the eight co-fractionation datasets by randomly separating each datasets’ replicates into two groups and taking each network as the union of unique edges from each group. After clustering each pair of networks, it was possible to calculate predicted reproducibility (*repJ*) and actual reproducibility between experimental replicates (*Ji*) for each cluster. Figure 8 shows that these values were correlated for all clustering algorithms. Thus, we find that *clust.perturb* can accurately predict which clusters will fail to be reproduced in a second experiment (Fig. 8). On the basis of this observation, we suggest that *clust.perturb* may be particularly useful for planning follow-up studies in situations where failure to reproduce clusters that were apparent in an initial experiment would be costly or time-consuming.

**Figure 8.**
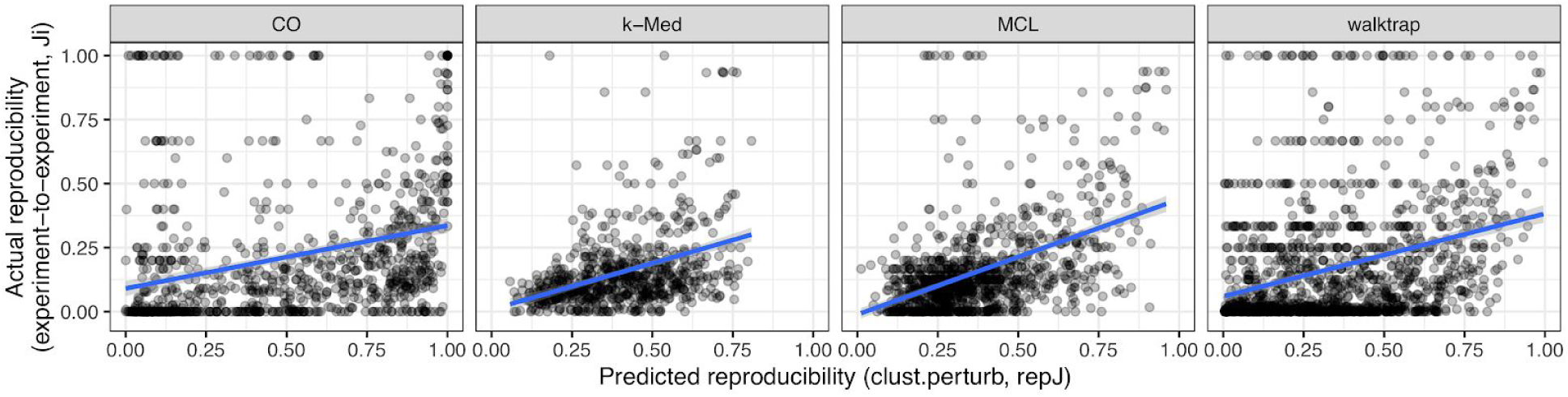
*clust.perturb* predicts real-world, experiment-to-experiment cluster reproducibility. Two cluster sets were constructed from two independent networks from the same experiment (replicates from the same experiment were randomly separated into two groups), and *Ji* was calculated between the sets, simulating real-world network variation (x-axis). The predicted reproducibility *repJ* was calculated for each cluster by *clust.perturb* (y-axis). Each dot represents a cluster. Blue lines show best linear fit.

## DISCUSSION

We find that graph-based clustering amplifies network errors. Indeed, this phenomenon is repeatedly observed across multiple common graph-based clustering algorithms. Consequently, in the presence of very minor random network-to-network variations (i.e. noise), clustering results tend to be poorly reproducible. Even small changes to a network, such as rewiring 1% of edges, lead to cluster results rearranging by more than 50%. This phenomenon represents more than network errors simply being propagated to the level of clusters. Instead, network alterations appear magnified by clustering, such that the number of altered cluster edges can be many times the number of altered network edges. Importantly, this variance in cluster sets is due to network noise and not inherent randomness in the clustering algorithms: for all algorithms here, cluster sets do not vary if the network is unchanged.

Therefore, studies that attempt to experimentally validate these clusters or otherwise include them in downstream analyses are vulnerable to this property, and may ultimately analyze spurious arrangements particular to one dataset, rather than clusters with general, real-world meaning. This point is important because networks, especially biological networks such as protein-protein interactomes, often contain errors in the range studied here (e.g. 10-50% false positives). If clustering accentuates these errors, then graph-based network clustering as a tool may be less useful to researchers than expected, at least in some contexts.

One explanation for this difficulty is that graph-based clustering is an inherently ill-posed problem (Jain 2010; Al-Razgan and Domeniconi 2006). That is, there are many sets of clusters that could be described by a given network. For example, a fully-connected network of three nodes and three edges could be a result of a three-member cluster, three two-member clusters, or any combination of them. Larger networks have even greater numbers of potential clusterings, especially in situations where subsets of clusters exist, such as the 40S and 60S subunits of the full 80S ribosome. This means that the solution space of clustering contains many solutions that are equally correct, and without more constraints it may be easy for noise to result in the selection of one over the others.

Our analysis also recapitulates previous observations that measuring the similarity of two sets of clusters can be ambiguous (Gates et al. 2017). Previous studies have looked at the robustness of clustering results to noise, particularly interactome clustering (Brohée and van Helden 2006; Sloutsky et al. 2013; Vlasblom and Wodak 2009; Freytag et al. 2018). Notably, Brohee et al. (2006) reported that clustering results appeared robust to both the addition and removal of interactions from the interactome. However, due to the ambiguity and difficulty with measuring the similarity between clustering sets, it is possible that these studies were, in fact, measuring something other than the “similarity” that we propose would correspond to a biologist’s intuition about how clusters should behave. Indeed, (Brohée and van Helden 2006) use geometric accuracy (GA) to measure similarity, a metric that we show can produce a score of 1 (i.e. perfect agreement) between non-identical cluster sets when set 2 contains proteins not found in set 1, a situation which is common.

To address some of the limitations of graph-based clustering discussed here, we developed *clust.perturb* (https://github.com/GregStacey/clust-perturb, https://rstacey.shinyapps.io/clust-perturb-tool/), a tool that provides metrics for cluster reproducibility (*repJ*) and the reproducibility of nodes within clusters (*fnode*). Because clusters fragment predictably in response to random noise (Fig. 5), it is possible to represent real, future alterations to the network (for instance, changes to a network across replicates of an experiment), with *in silico* network perturbations (that is, random rewiring of edges within the network). We found that cluster robustness in response to *in silico* network noise was predictive of clusters that were stable across networks derived from independent experiments (Fig. 8), and conversely, *clust.perturb* identified clusters that were spurious network associations that were not reproduced in subsequent experiments. Therefore, *clust.perturb* can provide additional computational evidence that identifies robust clusters, which are more likely to carry true, real-world meaning (Fig. 7).

Using simulated noise to predict the effects of future network alterations relies on noise being representative of those real-world alterations. In this study, we simulate network noise by the addition or removal of edges between existing nodes, i.e. the same nodes with altered edges. However, some real-world network alterations differ from this, for example in the case of subsequent CORUM releases, where much of the network differences result from the addition or removal of nodes (proteins). Future work remains to see how this type of noise (node addition and removal) affects cluster reproducibility.

## SUPPLEMENTARY FIGURES

**Supplementary Figure 1.**
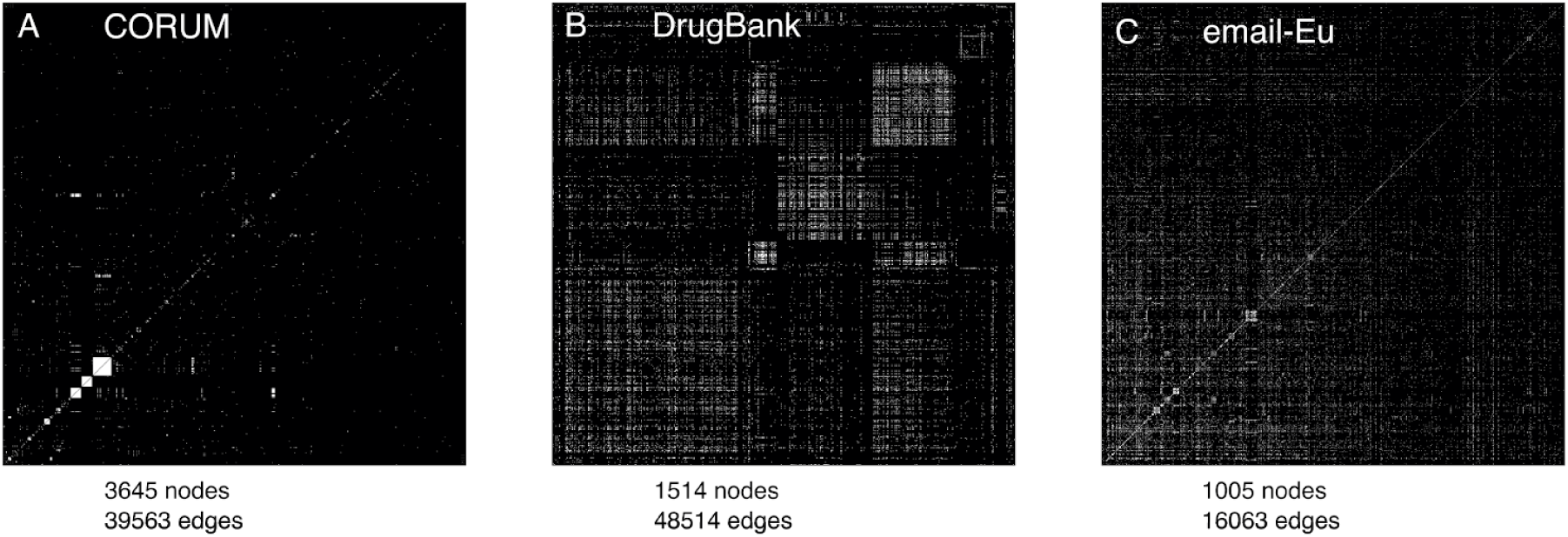
Adjacency matrices of the three network datasets. A) CORUM network with nodes (proteins) ordered by their first appearance in a CORUM complex. B) DrugBank network. Since there is no ground truth cluster set, nodes (drugs) are ordered using the optimal cluster assignment obtained during parameter optimization. C) email-Eu network with nodes (research institute members) ordered by faculty affiliation.

**Supplementary Figure 2.**
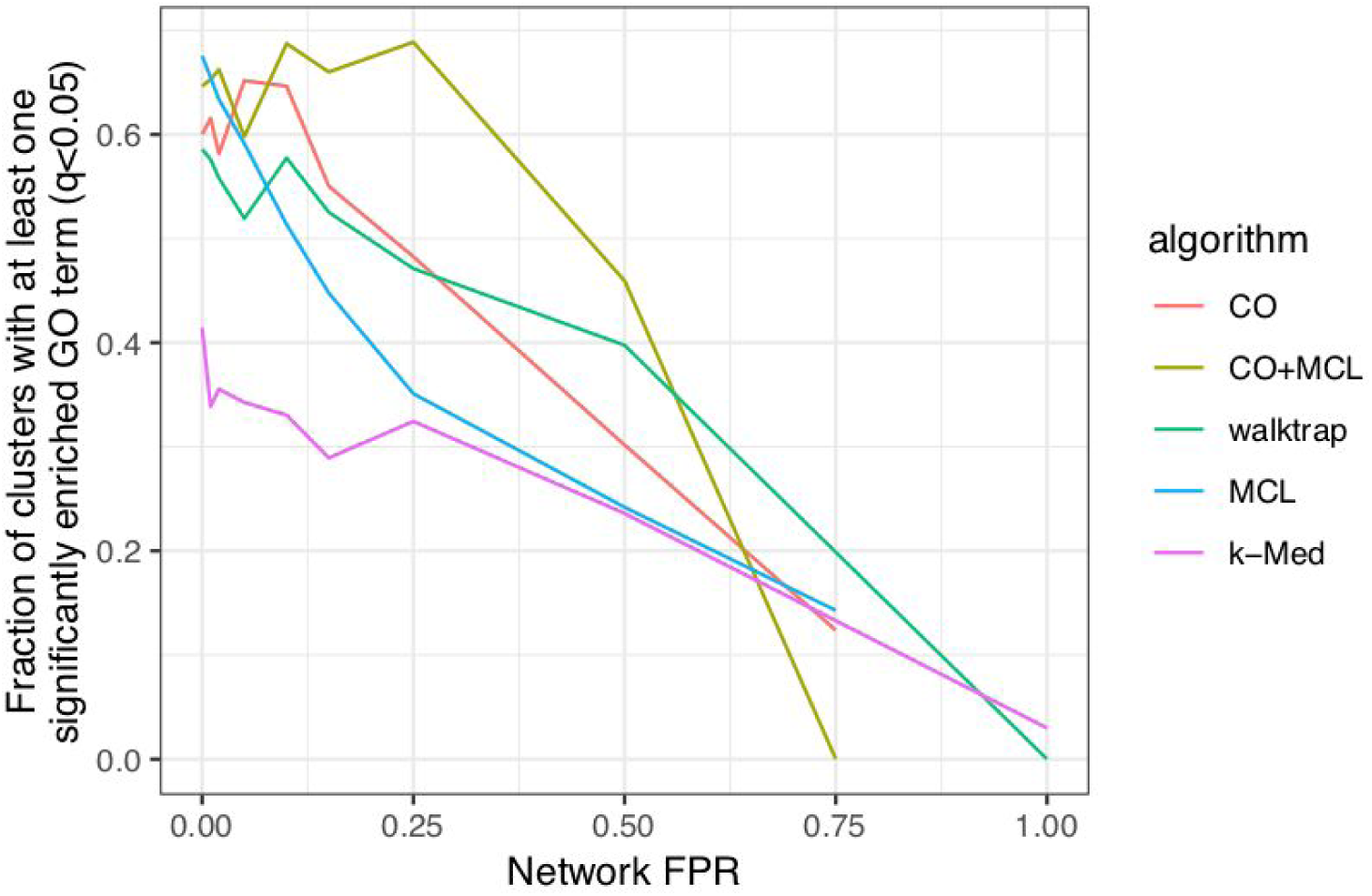
GO enrichment of clustered proteins decreases with network errors.

**Supplementary Figure 3.**
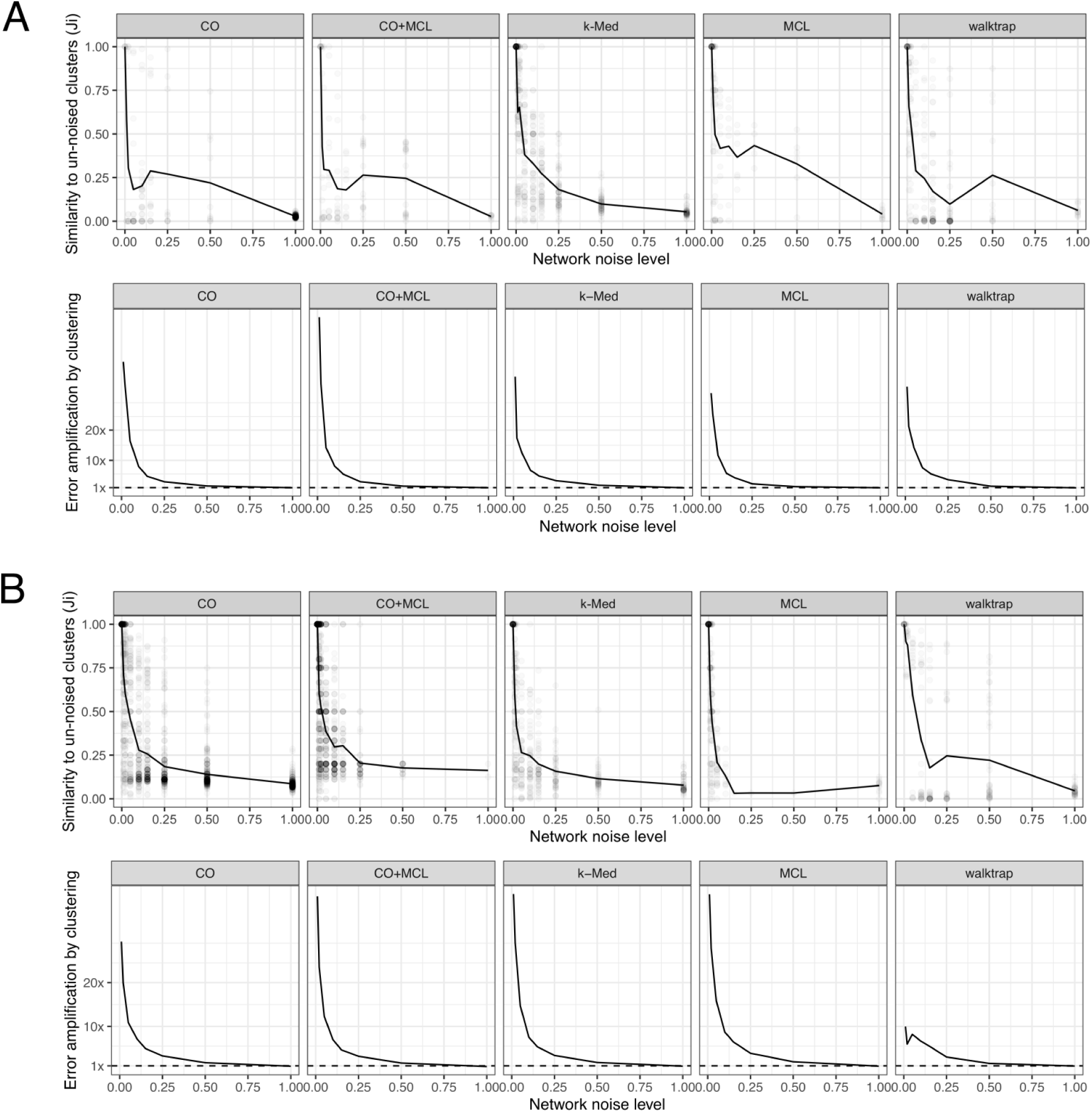
Clustering amplifies network noise (DrugBank and email-Eu). Complementary analysis to Figure 3 for network datasets DrugBank (A) and email-Eu (B).

**Supplementary Figure 4.**
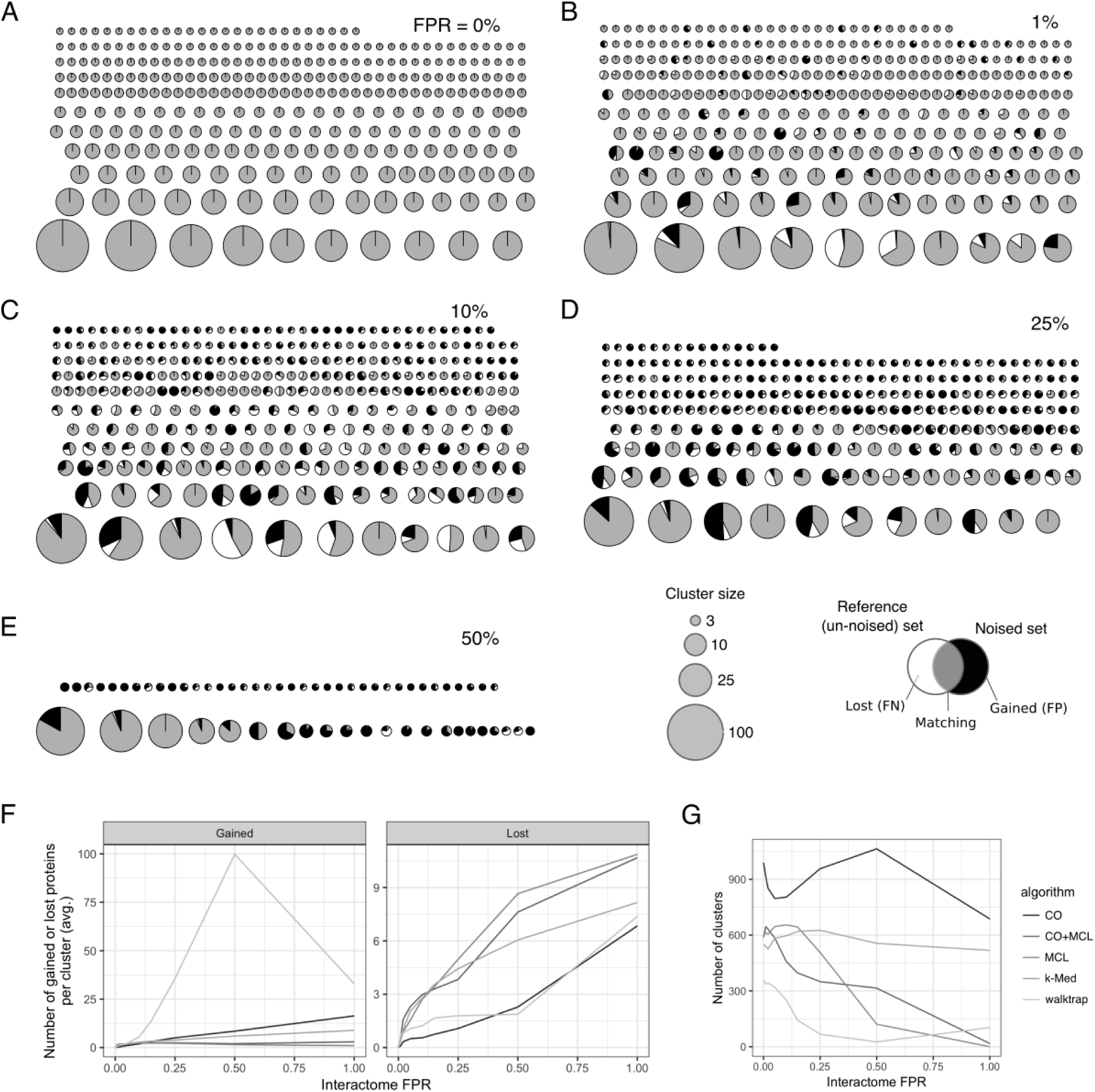
Rearrangement of clustering results in response to interactome noise - visualization and simple counting statistics. A-E) MCL clustering of the binarized CORUM network after addition of varying levels of interactome noise. Shading shows the agreement with clustering results from the original network. Grey shows overlap, white shows proteins that are lost after adding interactome noise, and black shows proteins that are added. F) Average numbers of gained and lost proteins per cluster. G) Total number of clusters in each clustering set.

**Supplementary Figure 5.**
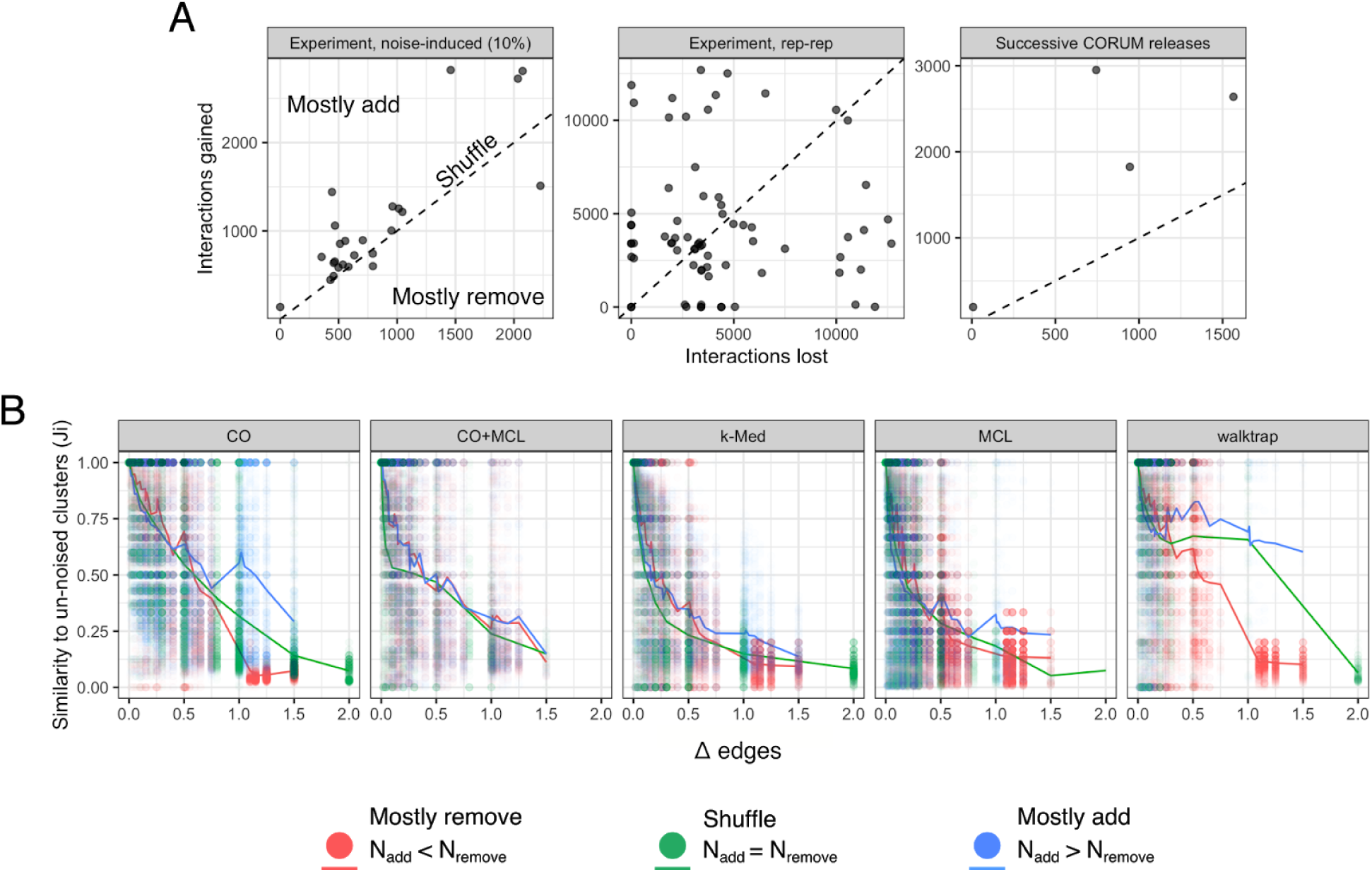
Edge addition and removal, effects on cluster reproducibility. A) Number of edges lost and gained (x- and y-axis) between different networks. Left: networks from the 28 experimental datasets compared to their networks after adding 10% noise (see Figure 4). Middle: pairs of replicates from the same experiment in the 28 experimental datasets. Right: loss and gain from one CORUM version to the subsequent CORUM version. B) Complementary analysis to Figure 3B, but including all combinations of edge removal and addition. Edges were added or removed in proportion to the original network size (0%, 1%, 2%, 5%, 10%, 15%, 25%, 50%, and 100%). 81 total combinations (9×9) of edge addition and removal. Δedges is the sum of the added and removed fractions, e.g. 50% removed and 25% added gives Δedges=0.75.

**Supplementary Figure 6.**
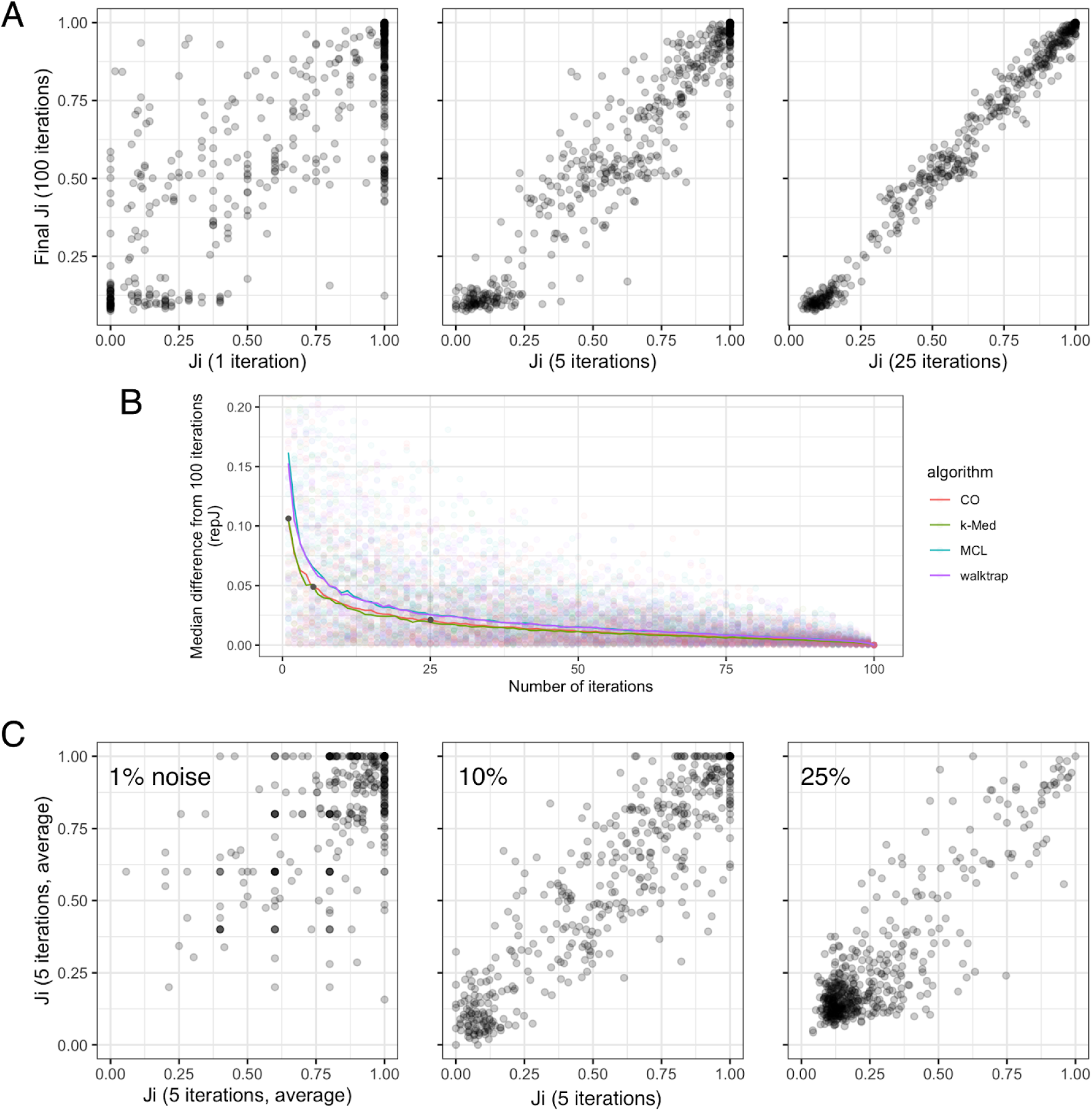
Effects of *clust.perturb* parameters. A) *Ji* converges towards a final value within few iterations. *Ji* from 1 noise iteration (left), 5 iterations (middle), and 25 iterations (right, x-axis) vs *Ji* from 100 iterations (y-axis). ClusterONE, 10% noise. B) Absolute difference from 100-iteration *Ji* as a function of number of iterations. Panels from A are shown by grey dots. C) Noise level should be chosen to adequately differentiate clusters, as in middle panel (10% noise). Left: too little noise, meaning most clusters remain unchanged (*Ji*=1). Right: too much noise, meaning most clusters are disrupted (*Ji* close to 0).

